# Combined Cartilage Thickness and Mechanical Property Mismatch Drives Local Strain Amplification at the Patellar Osteochondral Allograft Interface

**DOI:** 10.64898/2026.05.13.724923

**Authors:** Michael A. Hernández Lamberty, John A. Grant, Ellen M. Arruda, Rhima M. Coleman

**Affiliations:** Department of Mechanical Engineering, University of Michigan, Ann Arbor, Michigan, USA; Department of Biomedical Engineering, University of Michigan, Ann Arbor, Michigan, USA; Program in Macromolecular Science and Engineering, University of Michigan, Ann Arbor, Michigan, USA; Department of Orthopaedic Surgery, University of Michigan, Ann Arbor, Michigan, USA

**Keywords:** finite element analysis, cartilage, osteochondral allograft transplant, patella, patellofemoral joint

## Abstract

Patellar osteochondral allograft (OCA) transplantation is widely used to treat large full-thickness cartilage defects, yet long-term failure and reoperation rates remain high. Although surface congruity and osseous integration are emphasized clinically, cartilage thickness and mechanical compatibility between donor and recipient are not considered. Our previous work suggests that cartilage thickness mismatch can amplify local deformation at the graft boundary, potentially compromising graft longevity. This study investigates how combined mismatches in cartilage thickness and mechanical properties influence the local strain environment at the patellar OCA interface. Simplified two-dimensional axisymmetric finite element models of patellar OCA repair were developed in ABAQUS. Donor-to-recipient cartilage thickness ratios ranging from 0.33 to 3.25 were evaluated together with donor-recipient Young’s modulus mismatches (2.5-7.0 MPa). Cartilage was modeled using homogeneous linear elastic and functionally graded material formulations to account for depth-dependent stiffness. A compressive pressure of 1.0 MPa was applied to represent patellofemoral joint loading, and peak compressive and shear strains were quantified at the graft boundary. Cartilage thickness mismatch produced localized high-strain regions (HSR) of compressive and shear strain at the donor-recipient interface that were absent in thickness-matched constructs. Strain amplification increased with both thickness and mechanical property mismatch. Compressive strain exhibited directional asymmetry, with donor-side-thicker configurations producing greater amplification than recipient-side-thicker configurations. Incorporating depth-dependent cartilage stiffness reduced peak strain magnitudes but did not eliminate mismatch-driven strain amplification. These findings demonstrate that cartilage thickness and mechanical disparity can create HSR at the patellar OCA graft boundary that may predispose grafts to impaired integration and long-term failure.

## 1. Introduction

Articular cartilage is a specialized soft tissue found at the end of the bones of diarthrodial joints. It provides a near-frictionless surface for the load transfer between bones (Sophia Fox et al. 2009). This mechanical function is achieved through its complex hierarchical structure composed of a depth-dependent collagen fibril network. Overall, the articular cartilage is composed of an extracellular matrix (ECM) that consists primarily of water, collagen, and proteoglycans. Proteoglycans contain negatively charged glycosaminoglycan side chains that attract mobile ions and water into the tissue. The resulting influx of fluid generates osmotic swelling pressure that is resisted by the tensile stiffness of the collagen fibril network which enables articular cartilage to withstand compressive loads (Soltz and Ateshian 1998; Huang et al. 2001). Variations in the collagen fiber orientation, and proteoglycan and chondrocyte concentrations across the tissue depth give rise to the superficial, middle and deep zones of the articular cartilage, each with distinct structural organization and mechanical properties (Buckwalter et al. 1994; Antons et al. 2018).

Cartilage injuries are observed in up to 66% of patients undergoing knee arthroscopy procedures (Curl et al. 1997; Årøen et al. 2004). Full-thickness chondral injuries are observed in 4.6% - 6.2% of all patients, but the prevalence of full-thickness injuries increases to 36% in athletes undergoing knee arthroscopy (Flanigan et al. 2010). Among all the locations of chondral injuries that are found in the knee, approximately 20% are located in the patella (Curl et al. 1997). Cartilage has limited regenerative capacity due to its being both aneural and avascular. This often means that surgical methods are necessary to repair full-thickness injuries of the articular cartilage. If these chondral injuries are not addressed, they may compromise the joint function and accelerate the onset of osteoarthritis (Bhosale and Richardson 2008).

Osteochondral allograft (OCA) transplantation is the standard procedure for treating articular cartilage defects greater than 2 cm^2^ (Hevesi et al. 2021). The OCA transplant includes a cylindrical graft of healthy cartilage and the subchondral bone that is then press-fit into the recipient’s prepared defect site (Williams et al. 2004; Montgomery et al. 2014). One of the main benefits of the OCA procedure is that in a single-stage operation, the cartilage surface is restored. OCA has gained popularity among athletes with full-thickness defects due to its faster rehabilitation times and, consequently, faster return to play (Krych et al. 2016). Despite the advantages, OCA procedures are not without drawbacks. Previous work done by Familiari et al. showed that OCAs fail at a rate of 13% at 5 years, 21% at 10 years, 27% at 15 years, and 33% at 20 years following the procedure (Familiari et al. 2018). Patellar OCA transplant procedures have also shown high rates of failure and reoperation long term; at 15 years following the procedure, up to 20% of grafts failed, and 52% of the patients required reoperation (Chahla et al. 2019). The underlying mechanism behind OCA transplant failure in the patellofemoral joint remains poorly understood.

The primary parameters for successful OCA transplant procedures are articular cartilage congruity and osseous integration (Lai et al. 2022). However, both biological and mechanical factors influence the success of the procedure. In the patellofemoral joint, residual maltracking of the patella, joint instability, or incongruent subchondral surfaces can exacerbate the risk of OCA failure (Haber et al. 2019; Ackermann et al. 2019; Patel et al. 2021). Although the consequences of graft elevation and subchondral surface incongruity have been studied and are actively managed clinically, cartilage thickness mismatch between the donor and recipient tissue is not currently considered in graft selection protocols.

Our group previously showed that donor-to-recipient (D/R) cartilage thickness disparity resulting from OCA transplantation in the patella can lead to local stress concentrations up to twice the applied surface pressure near the graft boundary in the patella (Rosario et al. 2023). These high-strain regions (HSR) provide a potential mechanical mechanism for the failure of OCA transplants. Elevated stresses and strains at the graft boundary can have large biological consequences for the cartilage chondrocytes. Previous in vitro studies using cyclic compressive loading reported reduced chondrocyte viability at applied stresses of ∼6 MPa and above, with more severe loading procedure producing greater cell loss and matrix damage (Clements et al. 2001; Thibault et al. 2002). Other work using articular cartilage explants demonstrated that compressive strains exceeding ∼18% can induce chondrocyte apoptosis in the deep zone of the cartilage (Bartell et al. 2015; Huang et al. 2020). Notably, during daily activities such as running, patellofemoral contact pressures can reach ∼5 MPa with peaks approaching ∼10 MPa (Huberti and Hayes 1984; Lenhart et al. 2015). However, these in vivo contact pressures are transient and distributed, whereas observed in vitro injury thresholds depend on the testing protocol used, which makes it difficult to identify a physiological articular cartilage death threshold. When local strain amplification occurs at the donor-recipient interface of the OCA graft, physiological joint loads may generate localized strains that approach or exceed levels associated with chondrocyte injury in experimental studies. Thus, mechanical mismatches at the graft boundary are not only structural problems but also may trigger biological responses that contribute to progressive degeneration and graft failure.

In addition to geometric mismatch, mechanical property mismatch between the donor and recipient cartilage might lead to failure. Articular cartilage is an inhomogeneous tissue that has spatially varying mechanical properties (Deneweth et al. 2013, 2015; Antons et al. 2018). Prior studies have shown that the material constants for linear elastic, biphasic, or hyperelastic cartilage material models vary across joints and individuals (Armstrong and Mow 1982; Pierce et al. 2009; Antons et al. 2018). As a result, donor cartilage used for OCA transplantation may have a different stiffness than the surrounding recipient cartilage, even when the graft is geometrically well matched. This mechanical property disparity may alter load transfer across the donor-recipient cartilage interface. Therefore, the goal of this study was to identify the combined effect of D/R cartilage thickness and mechanical property mismatches on the local strain environment in a human patella OCA finite element (FE) model. We hypothesized that a larger disparity in D/R cartilage thickness compounded by a large disparity in the mechanical properties between the donor and recipient would produce higher magnitude strains in the HSR.

## 2. Methods

### 2.1 Finite Element Model

#### 2.1.1 Model Geometry

Simplified two-dimensional axisymmetric FE models of the patella cartilage with an osteochondral allograft (OCA) were created in ABAQUS 2022 (SIMULIA) to understand the effects of donor-recipient cartilage thickness mismatch and mechanical property mismatch. The model geometry and the donor-to-recipient (D/R) ratio were based on previously obtained nano-computed tomography (CT) measurements of patellar OCA transplants performed by a sports medicine orthopaedic surgeon on cadaveric patellae (Patel et al. 2021). The image acquisition and segmentation methods were previously described by Rosario et al. (Rosario et al. 2023).

Using the characteristics of the scans, the donor cartilage was modeled with a radius of 8 mm, while the recipient cartilage had a radius of 28 mm. The geometric mismatch was modeled as a sharp geometric transition at the donor-recipient interface. Cartilage thickness in the models was varied by changing the donor cartilage thickness while keeping the recipient cartilage thickness at 2 mm, resulting in donor-to-recipient (D/R) thickness ratios ranging from 0.33 to 3.25 (**Fig. 1**). These values were chosen based on the thickness values observed in the nano-CT scans provided for this study. Nine total D/R thickness ratios were modeled; four ratios in which the donor cartilage is thicker, four ratios in which the recipient cartilage is thicker, and a control ratio in which both cartilage sections have an equal thickness (**Table 1**).

**Fig 1:**
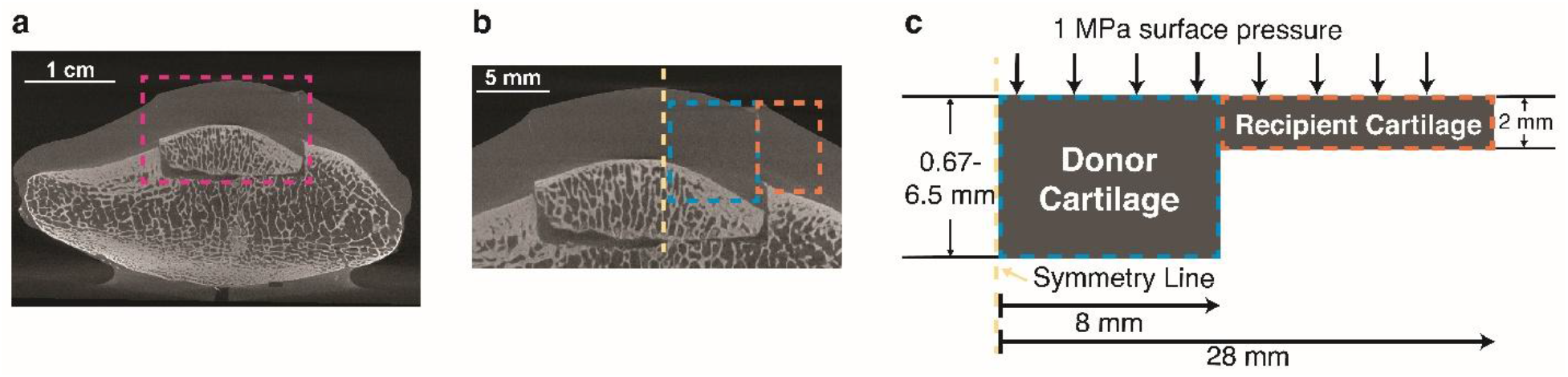
(a) Example nano-CT image of patella following a successful OCA procedure. In this image, cartilage thickness mismatches can be observed on both sides of the OCA graft. (b) The sections used to create the axisymmetric FE models in ABAQUS. The green line represents the line of symmetry for the model, the blue box represents the donor cartilage section, and the yellow box represents the recipient cartilage section. (c) Example of two-dimensional axisymmetric model.

**Table 1:** Nine total Donor-to-Recipient (D/R) thickness ratio FE models were developed to understand the role of thickness mismatch on the predicted compressive and shear strains at the graft interface. The control model was determined to be the FE model for which both the recipient and donor cartilage equal thickness. Four models were developed for which donor section was thicker than the recipient, and four models for which the recipient was thicker than the donor.

| Control Model | Thicker Donor Models<br>(D/R > 1) | Thicker Recipient Models<br>(D/R < 1) |
| --- | --- | --- |
| D/R = 1 | D/R = 1.25 | D/R = 0.80 |
|  | D/R = 1.50 | D/R = 0.67 |
|  | D/R = 2.00 | D/R = 0.50 |
|  | D/R = 3.25 | D/R = 0.33 |

#### 2.1.2 Mesh Generation

The FE models were discretized using 8-node biquadratic axisymmetric elements (CAX8). A quadratic mesh was generated with an average element edge length of approximately 50 μm throughout the whole cartilage. To increase the accuracy of the local strain gradients near the donor-recipient interface, the mesh was progressively refined within 2 mm of the graft boundary. The refinement linearly decreased the edge length of the quadratic elements from 50 μm to 5 μm at the interface. This refinement method was chosen to capture the localized high-strain region observed near the graft boundary. A mesh convergence analysis was performed by progressively refining the mesh near the graft boundary. Changes in the primary strain outcomes between successive mesh refinements were less than 3%, indicating convergence while avoiding a disproportionate increase in computation time.

#### 2.1.3 Material Models

##### 2.1.3.1 Homogenous Linear Elastic Models

In the initial set of FE analyses, the articular cartilage was modeled as a homogenous isotropic linear elastic material. Poisson’s ratio of 0.46 was used for all models (Jin and Lewis 2004). The stiffness or Young’s modulus was varied between 2.5 MPa and 7.0 MPa to represent reported ranges of human articular cartilage stiffness under physiological loading conditions and at various locations through the thickness of the cartilage (Chen et al. 2001; Antons et al. 2018).

Mechanical property mismatches were introduced by independently assigning Young’s modulus values to the donor and recipient cartilage regions. For each D/R thickness ratio, simulations were performed such that either the donor or the recipient cartilage Young’s modulus was held constant at 7.0 MPa while the opposing cartilage section was varied between 2.5 MPa and 7.0 MPa, resulting in a total of eleven mechanical property combinations (**Table 2**). A thickness-matched and mechanically matched model (D/R = 1; donor modulus = recipient modulus = 7.0 MPa) served as the control condition.

**Table 2:** Eleven total combinations of linear elastic mechanical properties (Donor Stiffness/Recipient Stiffness) were implemented as the material model used for the cartilage sections in the FE model to understand the role of mechanical property mismatch between and donor and recipient cartilage sections. The control model was determined as the FE model with equal stiffness in both donor and recipient cartilage sections. Five models were developed in which the donor cartilage was fixed at a stiffness of 7.0 MPa and the recipient cartilage section varied from 6 MPa to 2.5 MPa. Five other models were developed in which the recipient cartilage was fixed at a stiffness of 7.0 MPa and the donor cartilage varied from 6 MPa to 2.5 MPa.

| Control Model<br>(MPa/MPa) | Stiffer Donor Cartilage<br>(MPa/MPa) | Stiffer Recipient Models<br>(MPa/MPa) |
| --- | --- | --- |
| 7.0 / 7.0 | 7.0 / 6.0 | 6.0 / 7.0 |
|  | 7.0 / 5.0 | 5.0 / 7.0 |
|  | 7.0 / 4.0 | 4.0 / 7.0 |
|  | 7.0 / 3.0 | 3.0 / 7.0 |
|  | 7.0 / 2.5 | 2.5 / 7.0 |

##### 2.1.3.2 Functionally Graded Material Models

To account for the known depth-dependent mechanical behavior of articular cartilage, an additional set of FE models incorporated a functionally graded material (FGM) formulation. A custom user-defined field subroutine (USDFLD) was implemented in ABAQUS to assign spatially varying Young’s modulus values as a function of the through-thickness coordinates of the cartilage. The modulus was prescribed to vary linearly from a starting value of 2.5 MPa at the superficial tangential zone of the cartilage to a value of 7.0 MPa at the deep zone (**Fig. 2b)**. These values are consistent with previous experimental zonal variations in cartilage thickness (Töyräs et al. 2001; Jin and Lewis 2004; Antons et al. 2018). The variation of the material properties was controlled using the FE model coordinate system, resulting in evenly spaced stiffness gradients through the cartilage thickness. All other modeling assumptions, including geometry, thickness ratio, mesh density, boundary conditions, and loading, were identical to those used in the homogenous material model.

**Fig 2:**
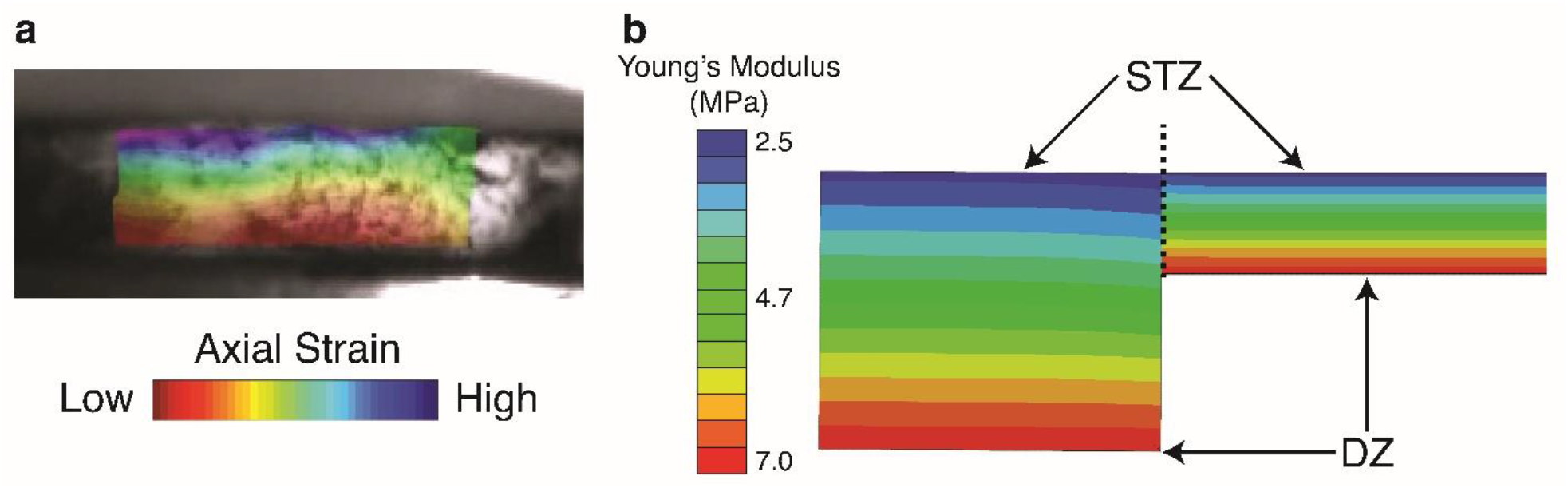
A functionally graded material (FGM) model was built in ABAQUS using a user-defined field (USDFLD) to define changing material properties along the thickness of the articular cartilage using literature-reported values. (a) Frame from a previous digital image correlation (DIC) experiment in our research group in which cartilage explant is compressed in an unconfined setup. The image demonstrates the change in axial strain through the thickness of the cartilage due to the changes in the mechanical properties of the articular cartilage. We used literature-reported values for the different zones of the cartilage to create the FGM model. (b) The developed USDFLD changes the Young’s modulus in equally spaced variations as a function of the coordinate system to achieve an FGM. The superficial tangential zone (STZ) was considered to have a stiffness of 2.5 MPa while the deep zone (DZ) of the cartilage was set to a stiffness of 7.0 MPa.

#### 2.1.4 Boundary Conditions and Loading

For both sets of material model simulations, the same boundary and loading conditions were implemented. The subchondral bone surface was modeled by setting all nodes at the bottom of the model to have fixed translation degrees of freedom. This boundary condition simulated the subchondral bone located beneath the articular cartilage. Axisymmetric boundary conditions were enforced along the central axis of the model. A uniform compression of 1.0 MPa was applied normally to the articular cartilage surface, which is within the range of physiological joint contact loading in the PFJ (Lenhart et al. 2015; Hart et al. 2022), and ensured model convergence. The compression was applied over a radius of 26 mm at the surface of the cartilage. A 2 mm distance was left from the edge of the cartilage to avoid any edge effects that might cause excessive distortion of the elements, which might cause the model not to converge.

### 2.2 Study Design and Data Analysis

Two parametric studies were conducted. In the first study, donor-recipient cartilage thickness ratio and mechanical property mismatch were varied using the homogenous linear elastic cartilage model described in Section 2.1.3.1. Nine D/R thickness ratios ranging from 0.33 to 3.25 were combined with eleven donor-recipient Young’s modulus combinations. The second study involved analyzing the effect of depth-dependent material behavior using the FGM material model described in Section 2.1.3.2. For this second study, the same set of D/R thickness ratios was evaluated while making sure the FGM varied through the thickness appropriately. This approach enabled us to understand the effect of cartilage thickness mismatch under more physiologically representative material assumptions.

The primary outcomes of interest were the minimum principal strain (compressive strain) and maximum shear strain. For each FE simulation, these quantities were extracted within 1 mm of either side of the donor-recipient graft boundary, where the HSR was observed (**Fig. 3a**). The peak values for each outcome were identified and compared across thickness and mechanical property combinations (**Fig. 3b**). For the homogenous linear material model, heatmaps summarizing the peak strain values were generated using a custom Python script to visualize trends associated with cartilage thickness and mechanical property mismatch. For the FGM, peak outcomes were compared directly to the extreme cases from the homogenous material simulation to assess the influence of depth-dependent material behavior.

**Fig 3:**
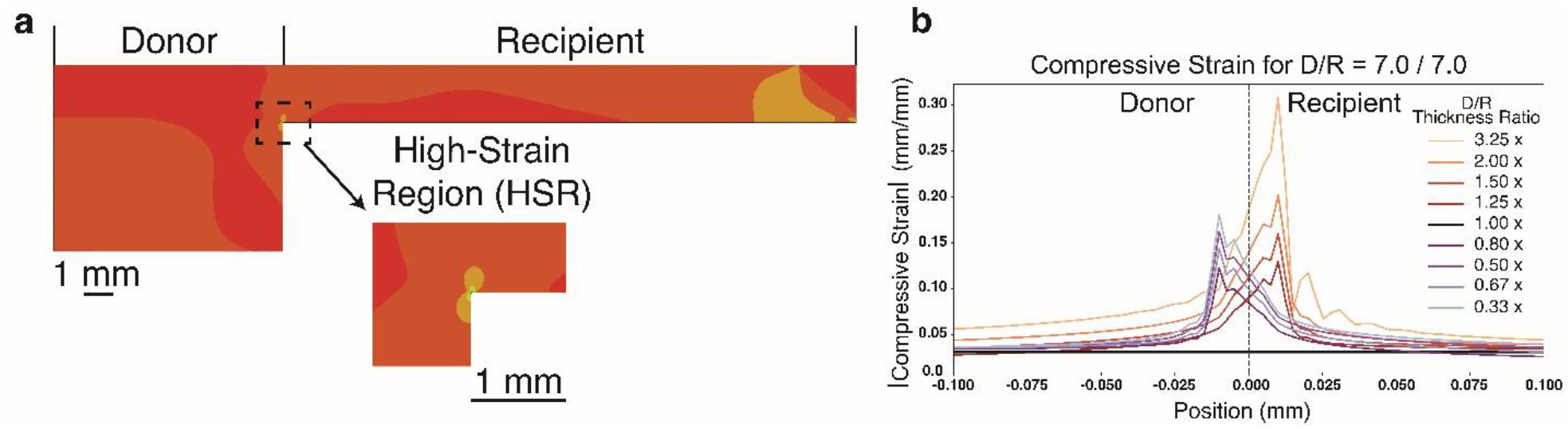
(a) Example image of ABAQUS FE model results for compressive strain. The high-strain region (HSR) is highlighted. The HSR is generated at the boundary between the donor and recipient cartilage of the OCA graft model. (b) Compressive strain for the control combination of mechanical properties (7.0/7.0 mechanical property combination). The limits of the x-axis of the plot are 0.1 mm in both directions from the OCA graft interface to emphasize the difference in compressive strain magnitude among thickness ratios.

## 3. Results

### 3.1 Impact of Cartilage Thickness and Mechanical Property Mismatch

Homogeneous linear elastic axisymmetric finite element models demonstrated that donor-recipient cartilage thickness mismatch produces localized HSR at the graft boundary that is absent in thickness-matched constructs. For all simulations, peak strains were localized near the donor-recipient interface and extended preferentially into the cartilage region that was thinner. For the control model, the peak compressive strain at the graft boundary was 0.03 (**Fig. 4a**). The shear strain at the graft boundary was negligible if either a cartilage thickness mismatch or a mechanical property mismatch was introduced into the model, localized regions of elevated compressive and shear strain developed at the donor-recipient interface, forming the HSR. Introducing a mechanical property mismatch while maintaining matched thickness (D/R = 1) resulted in increasing compressive strains at the interface. When the recipient cartilage was more compliant than the donor, in the most extreme case (7.0/2.5 MPa), peak compressive strain increased to 0.124, a 305% increase relative to the control. A smaller increase was observed when the recipient cartilage was stiffer than the donor (2.5/7.0 MPa) to a value of 0.121, which is 290% increase from the control. For the shear strain, increasing the mechanical property mismatch also produced increases in peak shear strain values, with the largest values again observed when the recipient cartilage was more compliant than the donor (7.0/2.5 MPa). In this case, the shear strain value increased to 0.148 (**Fig. 4b**). In the other extreme case of mechanical property mismatch, the shear strain value also increased to 0.129. This change in magnitude was smaller than the other combination but still substantial when compared to the control model.

**Fig 4:**
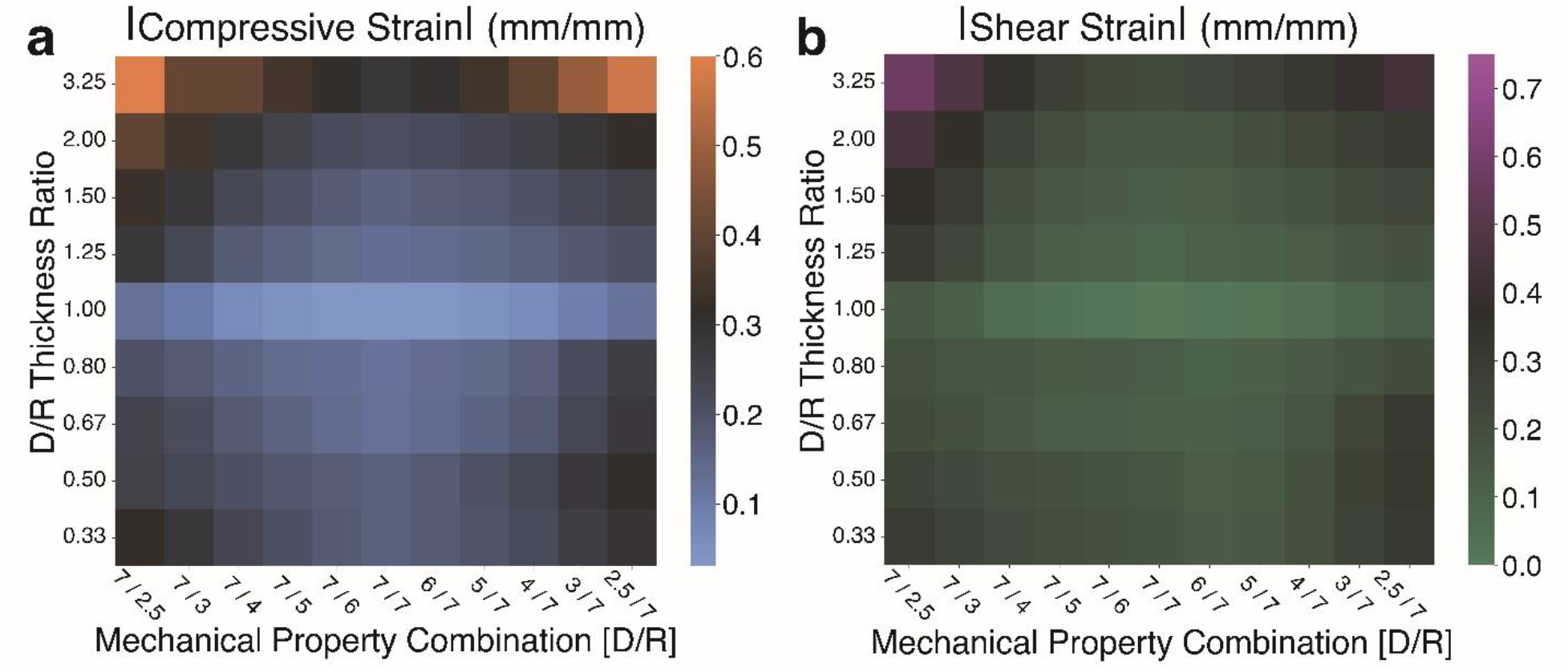
Heatmaps for the magnitude of all peak values of the outcomes of interest for the linear elastic material model. The compressive (a) and shear (b) strain heatmaps demonstrate a combinatorial effect in which an increase in disparity for both the D/R thickness ratio and mechanical property combination increases the magnitude of the strain at the OCA graft boundary.

Varying D/R cartilage thickness ratio while maintaining matched mechanical properties (7.0/7.0 MPa) also produced substantial strain amplification for both compressive and shear strains. Increasing the D/R thickness ratio from 1 to 3.25 increased the value of the peak compressive strain to 0.31, which demonstrates a 900% increase for the compressive strain. On the other hand, decreasing the ratio to 0.33 resulted in compressive strain peak values of 0.18, a 485% increase relative to control. For the shear strain, increasing the D/R thickness ratio from 1 to 3.25 increased the shear strain to 0.21, while decreasing the D/R thickness ratio from 1 to 0.33 resulted in shear strain of 0.17 from a value of 0 in the control model. For equivalent magnitudes of thickness mismatch, configurations with D/R > 1 consistently produced higher compressive strains than those with D/R < 1 (**Fig. 5**).

**Fig 5:**
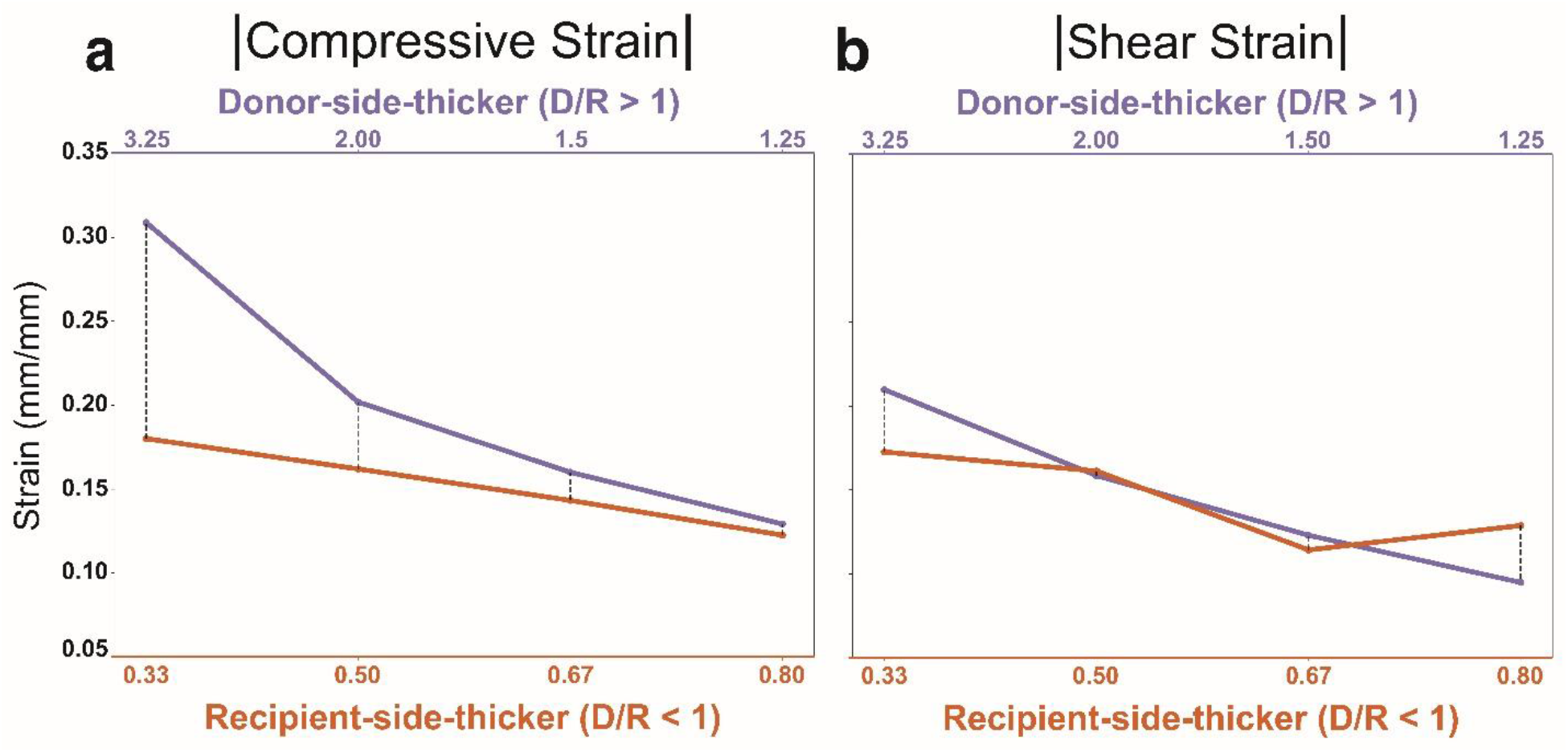
Comparison of peak compressive (a) and shear (b) strains at the OCA graft interface for the thickness-matched mechanical property model (7.0/7.0 MPa) across reciprocal cartilage thickness ratios. Compressive strain exhibited a consistent directional asymmetry, with donor-side-thicker configurations (D/R > 1) producing greater magnitudes than recipient-side-thicker configurations (D/R < 1) for all mismatch levels. In contrast, shear strain demonstrated magnitude-dependent directional behavior: donor-side-thicker models produced higher strain at the largest thickness disparity, whereas differences diminished or reversed as the thickness mismatch decreased.

When thickness mismatch and mechanical property mismatch were combined, a strong combinatorial effect was observed. The largest compressive strain was around 0.512 when a thick donor cartilage region (D/R = 3.25) was paired with a more compliant donor and stiffer recipient modulus combination (2.5/7.0 MPa), representing more than an order of magnitude increase relative to the control model strain of 0.03. For the shear strain, the largest strain value was 0.573, and it was observed when the D/R ratio was 3.25 and paired with the mechanical property combination of 7.0/2.5 MPa, where the donor cartilage was stiffer than the recipient cartilage.

### 3.2 Effect of Depth-Dependent Material Behavior on Strain

Peak compressive and shear strain values for the FGM models are shown in **Fig. 6** and are compared to the extrema cases for the homogenous linear elastic material FE model combinations (2.5/7.0 MPa and 7.0/2.5MPa). For the FGM control model (D/R = 1), peak compressive strain at the graft boundary was approximately 0.03, which was equal to the control value in homogenous models. Similarly, increasing cartilage thickness mismatch led to a monotonic increase in compressive strain for both D/R > 1 and D/R < 1 configurations. At D/R = 3.25, the peak compressive strain value increased to a magnitude of 0.284, while D/R = 0.33 resulted in a peak compressive strain of 0.172 (**Fig. 6a**), demonstrating increases of the compressive strain of 815% and 450%, respectively. Similar trends were observed for the shear strain. The control FGM model demonstrated a negligible shear strain at the graft boundary, whereas increasing the thickness mismatch produced larger shear strain magnitudes in the HSR. The shear strain magnitude had a value of 0.192 for the D/R = 3.25 model and 0.09 for the D/R = 0.33 model (**Fig. 6b**). Across all thickness ratios, FGM models produced lower peak strain magnitudes than the homogenous linear elastic models with extreme mechanical property mismatch.

**Fig 6:**
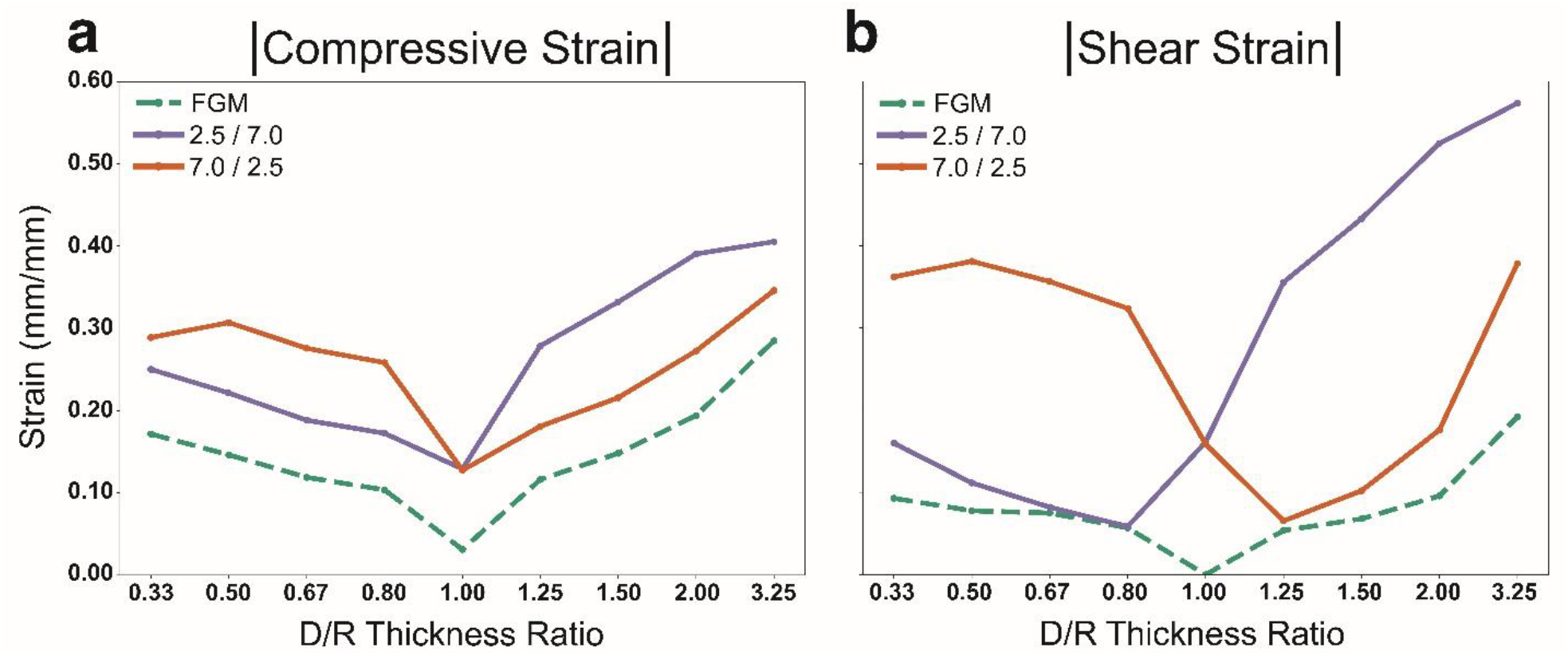
Comparison of compressive (a) and shear (b) strain magnitude between the FE models using the FGM material model and the extrema combinations of linear elastic material properties for donor and recipient cartilage sections. For both compressive and shear strain outcomes, the FGM models demonstrated lower strain magnitudes than the linear elastic cases. However, an increase in D/R thickness ratio did increase the magnitude of the strain outcomes for the FGM model, as in the linear elastic FE model.

## 4. Discussion

This study used simplified two-dimensional FE models to quantify how mismatches between donor-to-recipient cartilage thickness and mechanical property alter the local strain environment at the graft boundary of a patellar OCA transplant. Building on prior work demonstrating that thickness mismatch creates a localized amplification of strain at the graft boundary (Rosario et al. 2023), the present results demonstrate that these mismatches produce substantial amplification of both compressive and shear strains at the graft boundary. Importantly, these elevated strain environments persist across a wide range of thickness ratios and mechanical property combinations and remain present even when depth-dependent cartilage stiffness is incorporated. Together, these findings identify cartilage thickness and stiffness mismatch as mechanically robust contributors to localized strain amplification that may contribute to impaired graft integration and long-term viability of the patella OCA transplants.

The increase in the observed strain is interpreted as a thickness-driven strain concentration mechanism that is caused by the discontinuity in the cartilage thickness between donor and recipient sections and the change of stiffness across the donor-recipient interface. In matched constructions (D/R = 1), strain values are low or negligible, and they are distributed smoothly through the cartilage. In contrast, a thickness or stiffness mismatch in the model introduces a local constraint that redirects deformation toward the interface, resulting in the local HSR being observed. When a stiffness mismatch is present between the donor and recipient cartilage, the predicted strain increases further, particularly when stiffness differences coincide with a geometric mismatch. In these cases, thickness and stiffness driven constraints act synergistically to restrict deformation pathways on one of the sides of the graft interface, causing larger strains to develop. Importantly, these interactions are most pronounced for shear strain. This is consistent with prior experimental and computational studies demonstrating that shear deformation in articular cartilage is sensitive to surface integrity, collagen organization, and material heterogeneity (Maier et al. 2017; Trevino et al. 2017).

The results demonstrate that compressive and shear strains respond differently to cartilage thickness mismatches. Compressive strain exhibited a clear directional asymmetry, with donor-side-thicker configurations (D/R > 1) consistently producing greater magnitudes than recipient-side-thicker configurations (D/R < 1) across all reciprocal mismatch pairs. In contrast, shear strain displayed a magnitude-dependent directional response. While donor-side-thicker configurations generated larger shear strain at the highest mismatch levels, the differences diminished or reversed at smaller mismatch magnitudes. This suggests that compressive strain amplification is more uniformly sensitive to thickness asymmetry, whereas shear strain redistribution depends more strongly on the degree of geometric disparity at the graft boundary.

Incorporating depth-dependent cartilage stiffness through the FGM model definition reduced the peak strain magnitudes relative to the most extreme cases from the homogenous linear elastic FE models. However, the FGM FE model did not eliminate the strain amplification caused by the cartilage thickness disparities between the donor and recipient. The fact that the compressive and shear strain values increase as well when an FGM was used to describe the cartilage indicated that thickness-driven strain redistribution is not an artifact of uniform material assumptions, but rather a consequence of underlying geometry at the donor-recipient graft boundary. It is important to point out that the mitigating effect of the depth-dependent stiffness was more pronounced in the shear strain values observed in thicker donor configurations, where homogenous models exhibited the largest shear strain increase. For the thicker recipient, the differences in shear strain between FGM and homogeneous models were smaller but remained evident. Together, these findings indicate that while depth-dependent material behavior can decrease predicted peak strain magnitudes, it does not fundamentally alter the dominant role of cartilage thickness mismatch in shaping the local strain environment in the donor-recipient interface of the OCA graft.

From a biological perspective, elevated compressive and shear strains have been associated with chondrocyte injury (Bartell et al. 2015), apoptosis (Thomas et al. 2011), and matrix degradation (Trevino et al. 2017) in articular cartilage, particularly when repeatedly experiencing physiological joint loading. Within this framework, the localized HSR predicted in this study represents a mechanically plausible environment that could compromise chondrocyte viability and lateral integration at the graft boundary over time. The sensitivity of shear strain values to cartilage thickness disparity is especially important, as shear deformation has been noted as a driver of cartilage damage and cell death (Trevino et al. 2017; Ayala et al. 2021). Importantly, recent experimental work has demonstrated the feasibility of directly measuring full volume cartilage strain fields following OCA transplantation using displacement-encoded magnetic resonance imaging (Hernández Lamberty et al. 2026). The measurements revealed localized alteration in compressive and shear strain near portions of the graft rim under controlled indentation, providing experimental support for the existence of strain heterogeneity near the donor-recipient interface. Although the experiments were performed under sub-physiological loading, those results suggest that the HSR identified here are not purely theoretical and may manifest in vivo under functional joint loading.

Clinically, these findings suggest that cartilage thickness and stiffness mismatches can create localized mechanical environments that are not apparent just from surface congruity or osseous integration alone. Over time, repeated exposure to elevated compressive and shear strains at the OCA graft boundary may weaken the cartilage matrix, reduce chondrocyte viability, and predispose to degeneration of the cartilage at donor-recipient interface (Wilson et al. 2006; Bartell et al. 2015). This is especially true in the patellofemoral joint, where contact pressure of the patella cartilage can be 5, 7, or even 10 – 15 MPa for daily activities such as walking, stair ascent, or running (Huberti and Hayes 1984; Lenhart et al. 2015; Hart et al. 2022; Liao et al. 2018; Brechter and Powers 2002). Together, these results support consideration of cartilage-level compatibility during graft selection and surgical planning and highlight local HSR as a mechanistically meaningful indicator of graft vulnerability.

Several limitations should be considered when interpreting these findings. Firstly, cartilage was modeled using linear elastic material behavior, which does not capture viscoelasticity, anisotropy, poroelastic fluid pressurization, or time-dependent effects known to influence cartilage biomechanics (Huang et al. 2001; Lawless et al. 2017). However, the consistent strain amplification across homogenous and FGM formulations suggests that the primary conclusions are driven by thickness-dependent strain redistribution rather than by the specific constitutive models used. Additionally, the simplified axisymmetric geometry and loading conditions do not represent patient-specific anatomy, patellofemoral joint kinematics, or variations in contact across knee flexion angles. The models also assumed perfect lateral integration between donor and recipient cartilage and did not incorporate frictional contact, interfacial sliding, or incomplete osseous integration. Finally, the material properties were selected from literature-reported ranges rather than subject-specific measurements, and post-transplant tissue remodeling was not considered.

In summary, this study demonstrates that cartilage thickness mismatch in patellar OCA transplantation produces localized high-strain regions characterized by elevated compressive and shear strains at the donor-recipient interface through thickness-driven strain redistribution. Mechanical property mismatch further exacerbates these effects, while incorporating a depth-dependent cartilage behavior reduces but does not eliminate strain increase at the OCA graft interface. Shear strain exhibits strong sensitivity to cartilage thickness disparity and combined geometric-material mismatch. These findings indicate that donor-recipient cartilage thickness mismatch and mechanical incompatibility can create a HSR at the OCA graft boundary, providing a plausible mechanical pathway linking graft selection to impaired integration and eventual graft failure.

## Acknowledgements

The authors would like to thank Dr. Carla Nathaly Villacís Núñez, Dr. Ryan Rosario, and Peter Kuetzing for their feedback on model analysis and results.

## 5. Declarations

### 5.3 Data Availability

The finite element model files and derived data presented in this study are available from the corresponding author upon reasonable request.

### 5.4 Author Contributions

**Michael A. Hernández Lamberty:** Writing – original draft, Writing – review & editing, Conceptualization, Data curation, Formal Analysis, Investigation, Methodology, Software, Validation, Visualization. **John A. Grant:** Writing – review & editing. **Ellen M. Arruda:** Writing – review & editing, Conceptualization, Funding acquisition, Project administration, Resources, Supervision. **Rhima M. Coleman:** Writing – review & editing, Conceptualization, Funding acquisition, Project administration, Resources, Supervision.

